# A Genetic Screen Links the Disease-Associated Nab2 RNA-Binding Protein to the Planar Cell Polarity Pathway in *Drosophila melanogaster*

**DOI:** 10.1101/2019.12.23.887257

**Authors:** Wei-Hsuan Lee, Edwin Corgiat, J. Christopher Rounds, Zenyth Shepherd, Anita H. Corbett, Kenneth H. Moberg

## Abstract

Mutations in the gene encoding the ubiquitously expressed RNA-binding protein ZC3H14 result in a non-syndromic form of autosomal recessive intellectual disability. Studies in *Drosophila* have defined roles for the ZC3H14 ortholog, Nab2 (aka *Drosophila* Nab2 or dNab2), in axon guidance and memory due in part to interaction with a second RNA-binding protein, the fly Fragile X homolog Fmr1, and coregulation of shared Nab2-Fmr1 target mRNAs. Despite these advances, neurodevelopmental pathways regulated by Nab2 remain poorly defined. Structural defects in *Nab2* null brains resemble defects observed upon disruption of the planar cell polarity (PCP) pathway, which regulates planar orientation of static and motile cells. A kinked bristle phenotype in surviving *Nab2* mutant adults additionally suggests a defect in F-actin polymerization and bundling, which is also a PCP-regulated processes. To test for Nab2-PCP genetic interactions, a collection of PCP loss-of-function alleles was screened for modification of a rough-eye phenotype produced by Nab2 overexpression in the eye (*GMR-Nab2*) and subsequently for modification of *Nab2* null phenotypes. Multiple PCP alleles dominantly modify *GMR-Nab2* eye roughening and a subset of these alleles also rescue low survival and thoracic bristle kinking in *Nab2* zygotic nulls. Moreover, alleles of two X-linked PCP factors, *dishevelled* (*dsh*) and β *amyloid protein precursor-like* (*Appl*), rescue *GMR-Nab2* eye roughening in male progeny derived from hemizygous *dsh* or *Appl* mutant fathers, suggesting an additional effect inherited through the male germline. These findings demonstrate a consistent pattern of Nab2-PCP genetic interactions that suggest molecular links between Nab2 and the PCP pathway in the developing eye, wing and germline.

## INTRODUCTION

Mutations in genes that encode RNA-binding proteins often lead to tissue-specific disease pathology, particularly within the brain and nervous system (reviewed in Castello *et al*. 2013). Inactivating mutations in the human *ZC3H14* gene, which encodes a ubiquitously expressed zinc-finger, poly(A) RNA-binding protein, are linked to a monogenic form of non-syndromic autosomal recessive intellectual disability (reviewed in Fasken *et al*. 2019). The ZC3H14 protein and its homologs in the budding yeast *S. cerevisiae* and fruit fly *Drosophila melanogaster* (Nab2 in both species) interact with A-rich motifs in RNAs and restrict the length of mRNA polyadenosine (poly(A)) tails. Phenotypes associated with loss of Nab2/ZC3H14 are thus predicted to arise due to defects in post-transcriptional control mechanisms that involve poly(A)-dependent mRNA processing and nuclear export, stability, localization, and/or translation.

Studies in *Drosophila* indicate that Nab2/ZC3H14 share a conserved and necessary function in brain neurons (Pak *et al*. 2011; Kelly *et al*. 2014). Neuron-specific *Nab2* RNAi is sufficient to recapitulate phenotypic effects of *Nab2* loss, and neuron-restricted expression of either fly Nab2 or human ZC3H14 rescues developmental defects that otherwise occur in *Nab2* zygotic null animals. Notably, RNAi knockdown of Nab2 in motor neurons has no effect on survival or behavior (Kelly *et al*. 2014), implying a requirement for Nab2 in central nervous system (CNS) neurons. Consistent with this hypothesis, loss of Nab2 alters brain structure and impairs olfactory and courtship memory in *Drosophila*, while *ZC3H14* loss in mice alters hippocampal morphology and decreases working memory (Kelly *et al*. 2016; Rha *et al*. 2017; Collins *et al*. 2019). Mice lacking ZC3H14 have enlarged lateral ventricles and exhibit defects in a water-maze test of working memory, while Nab2-deficient flies exhibit defective axon guidance in the αβ lobes of the mushroom body (MB) and defective memory in courtship and aversive odor paradigms (Bienkowski *et al*. 2017). Within *Drosophila* brain neurons, Nab2 protein concentrates in the nucleus but is also found in cytoplasmic messenger ribonucleoprotein particles (mRNPs) in a physical complex with the fly Fragile-X protein Fmr1 (Bienkowski *et al*. 2017). This Nab2-Fmr1 complex is independent of linking RNA and correlates with translational repression of shared target mRNAs, e.g. *CaMkII* mRNA. Murine ZC3H14 co-sediments with a puromycin-sensitive 80S ribosomal fraction and localizes to developing axons and dendritic spines; in the latter compartment, ZC3H14 colocalizes with the post-synaptic guanylate kinase PSD95. In aggregate, these data indicate that cytoplasmic Nab2/ZC3H14 is found in neuronal mRNPs and likely plays roles in pre- and post-synaptic expression of mRNAs involved in axonogenesis and memory.

The majority of Nab2-regulated mRNAs are undefined, but similarities between phenotypes of a *Nab2* null allele and other mutants could identify candidate Nab2-regulated pathways. One such candidate is the Wnt/PCP (planar cell polarity) pathway, which controls the planar orientation of cells via localized effects on the F-actin cytoskeleton (reviewed in Yang and Mlodzik 2015). PCP involves two apically localized transmembrane complexes, Starry Night (also Flamingo)-Van Gogh-Prickle (Stan-Vang-Pk) and Starry night-Frizzled-Dishevelled-Diego (Stan-Fz-Dsh-Dgo), that interact across cell:cell junctions but are mutually antagonistic within cells, resulting in a polarized pattern of complex accumulation that propagates across the apical plane of an epithelium. PCP contributes to a number of developmentally programmed processes, including convergent extension, neural tube closure, and proximal-distal hair cell orientation (Yang and Mlodzik 2015). Significantly, the PCP factors are also involved in axon extension and guidance in the developing nervous systems of mouse, chicken, *Drosophila* and *C. elegans* (Sato *et al*. 2006; Srahna *et al*. 2006; Inaki *et al*. 2007; Hollis and Zou 2012; Ng 2012; Vandewalle *et al*. 2013; Ackley 2014; Gombos *et al*. 2015; Aviles and Stoeckli 2016). In *Drosophila*, loss-of-function alleles of the PCP factors *dishevelled* (*dsh*), *prickle* (*pk*), *frizzled* (*fz*) and *Van Gogh* (*vang*; also *strabismus, stbm*) disrupt axon projection into α and β lobes of the brain mushroom bodies (MBs) in a manner that resembles αβ MB defects in *Nab2* mutants (Ng 2012; Kelly *et al*. 2016). Nab2 loss also produces a prominent kinking defect in adult thorcic bristles (Pak *et al*. 2011), which are generated through the PCP-regulated process of actin polymerization and bundling (Yang and Mlodzik 2015). These phenotypic similarities between the effects of PCP and *Nab2* alleles on MBs and thoracic bristles suggest a potential link between post-transcriptional effects of Nab2 and PCP activity.

To use genetic means to test a Nab2-PCP link *in vivo*, a panel of alleles corresponding to PCP and PCP-related factors was screened for dominant modification of a rough-eye phenotype produced by eye-specific *Nab2* overexpression (*GMR-Nab2*) (Pak *et al*. 2011). This analysis reveals a consistent pattern of genetic interaction between PCP factors and exogenous *GMR-Nab2* that could also be observed with a null allele of endogenous *Nab2* (*Nab2*^*ex3*^). Nab2-PCP genetic interactions are apparent at multiple levels, including overall organismal viability, thoracic bristle morphology and patterning of the eye epithelium, and further supported by misorientation of wing hairs in *Nab2* mutants, a hallmark feature of most PCP factors in *Drosophila* (reviewed in Yang and Mlodzik 2015). Somewhat surprisingly, the modifying effect of *dishevelled* (*dsh*) and *β-Amyloid protein precursor-like* (*Appl*) alleles on *Nab2* gain (*GMR-Nab2*) or loss (*Nab2*^*ex3*^) can also be transmitted through the male germline independent of their inheritance, suggesting an additional, potentially epigenetic, link between PCP and Nab2.

## MATERIALS AND METHODS

### *DROSOPHILA* GENETICS

Crosses were maintained in 25°C humidified incubators with 12hr light-dark cycles (Shel•Lab). The *Nab2* alleles *ex3* (null), *pex41* (*precise excision 41* control), and *Nab2*^*EP3716*^ used for Gal4-driven *Nab2* expression, have been described previously (Pak *et al*. 2011). Lines obtained from the Bloomington *Drosophila* Stock Center (BDSC): *GMR-Gal4 dsh*^*a3*^, *dsh*^*6*^, *dsh*^*1*^, *Appl*^*d*^, *puc*^*MI11060*^, *pk*^*sple-13*^, *pk*^*sple-14*^, *pk*^*30*^, *wnt4*^*C1*^, *tap*^*MI10541*^, *fz*^*Jb*^, *vang*^*6*^, *stan*^*frz3*^, and *pygo*^*s123*^. Additional PCP lines were a gift of G. Pierre-Louis (present address Valencia College, FL) and J. Axelrod.

### IMAGE and DATA COLLECTION

Adult flies were imaged with a Leica DFC500 digital camera under white light. Images were processed with ImageJ and Adobe Photoshop to standardize brightness and convert to greyscale. Eye phenotypes were categorized as ‘Enhancer (Enh)’, ‘Suppressor (Sup)’, or ‘no effect’ based on visual assessment of pigmentation loss and disorganization of the retinal honeycomb structure (see **Table 1**). For survival assays, the hatching rate of non-Tubby pupae collected from a *Nab2*^*ex3*^*/TM6B*^*Tb,Hu*^ stock, either alone or also carrying the indicated PCP alleles (as in **Figure 2**), was determined by manual counting. To assess bristle kinking, humeral and scutal macrochaeta on the thorax were visually examined for the sharp kinks characteristic of *Nab2*^*ex3*^ adults (Pak *et al*. 2011). Adults with at least one kinked thoracic bristle were scored ‘positive’ for the kinking phenotype.

**Table 1.** *GMR-Nab2* modification by Wg/PCP alleles.

| modifier | phenotypic effect |  |
| --- | --- | --- |
|  | pigment | structure |
| *dsh*<sup>A3</sup> | Sup | Sup |
| *dsh*<sup>6</sup> | no effect | no effect |
| *Appl*<sup>d</sup> | Sup | Sup |
| *fz*<sup>JB</sup> | Sup | Sup |
| *Wnt4*<sup>c1</sup> | Enh | Enh |
| *pk*<sup>pk-sple13</sup> | no effect | no effect |
| *pk*<sup>pk-sple14</sup> | Sup | Sup |
| *pk*<sup>30</sup> | Sup | Sup |
| *puc*<sup>MI11060</sup> | Sup | Sup |
| *tap*<sup>MI10541</sup> | Sup | Sup |
| *stan*<sup>frz3</sup> | Sup | Sup |
| *vang*<sup>6</sup> | slight Sup | no effect |
| *pygo*<sup>s123</sup> | slight Enh | no effect |

\*Sup=suppressed \*Enh=enhanced

**Figure 1.**
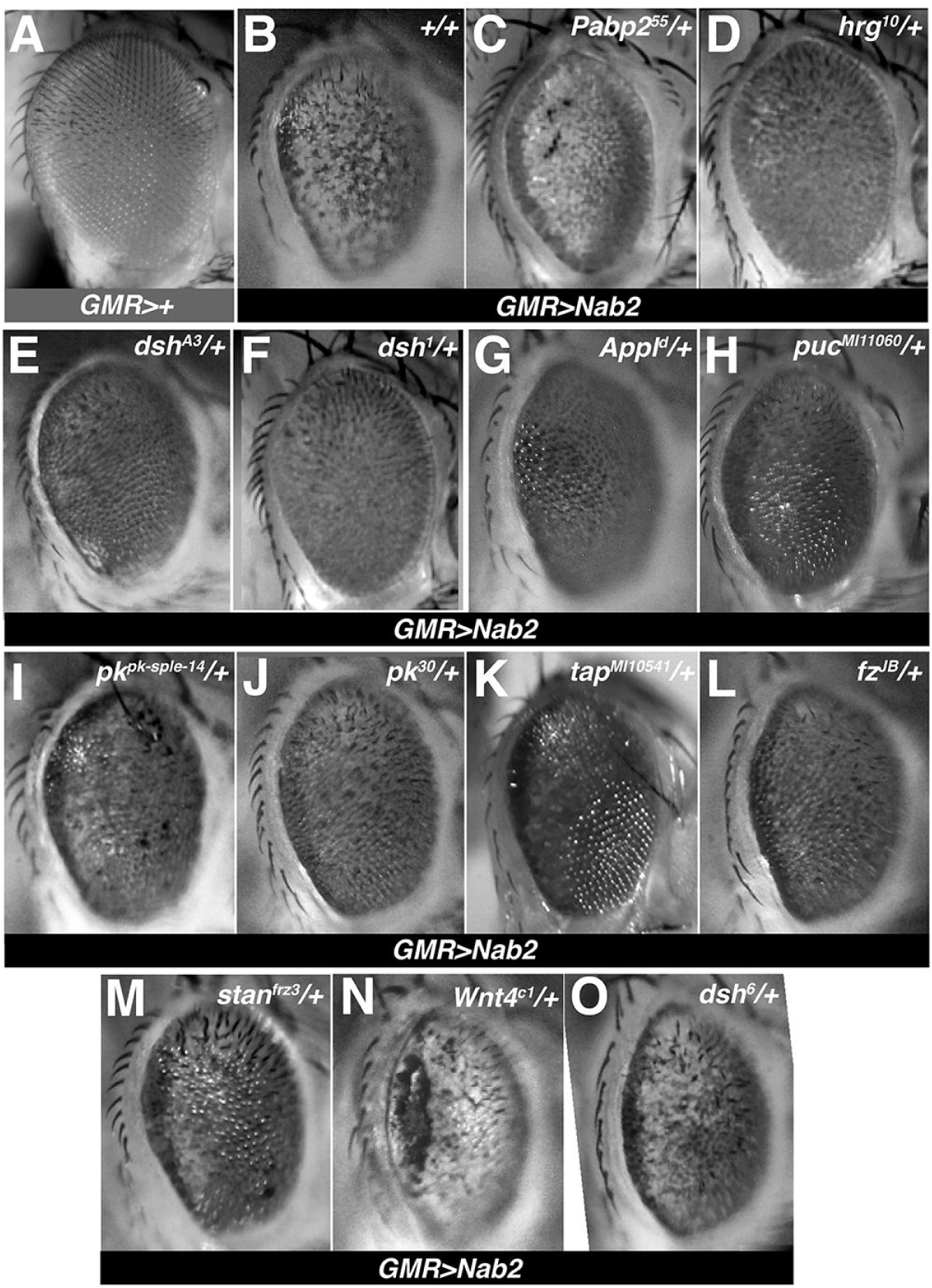
*GMR-Nab2* eye modification by Wg/PCP alleles. Images of (**A**) control, (**B**) *GMR-Nab2* and (**C-O**) *GMR-Nab2* adult female eyes also heterozygous for the indicated alleles. The images of *GMR-Nab2* eyes combined with the *Pabp2*^*55*^ (**C**) or *hrg*^*10*^ (**D**) loss-of-function alleles are provided as positive controls for enhancement and suppression (as in Pak *et al*. 2011).

**Figure 2.**
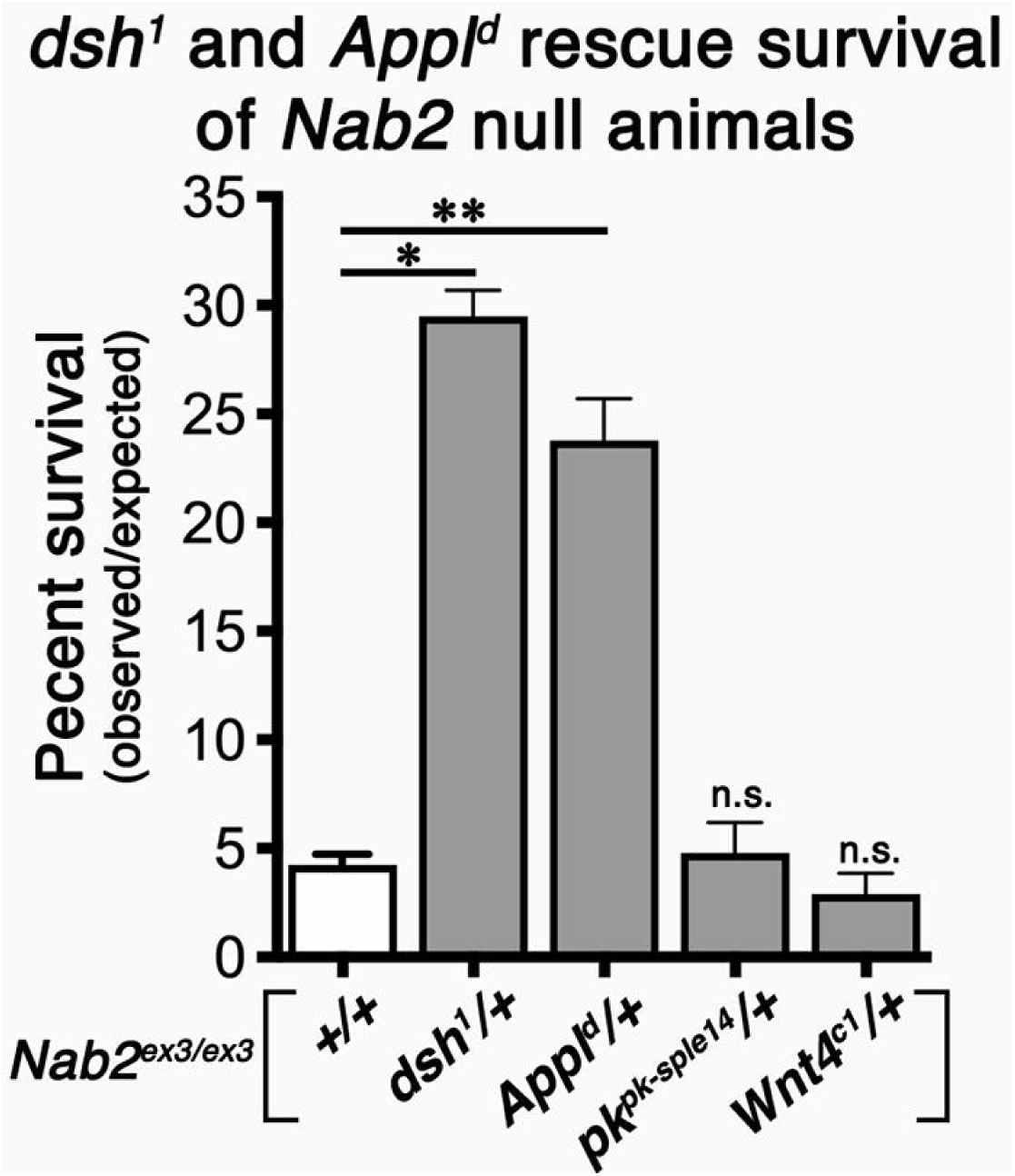
Modification of *Nab2*^*ex3*^ survival by a panel of four Wg/PCP alleles. Quantification of pupal eclosion rates among *Nab2*^*ex3*^ homozygous females (white fill), or *Nab2*^*ex3*^ homozygotes that are also heterozygous for *dsh*^*1*^, *Appl*^*d*^, *pk*^*pk-sple-14*^ or *Wnt4*^*c1*^. Data are presented as the number of flies that eclose (observed) vs the total number of pupae tracked (expected) from 3 separate crosses. Statistical significance is indicated (* p=0.0002; **p=0.001; n.s. = not significant).

### STATISTICAL ANALYSIS

Modifying effects of PCP alleles on *Nab2*^*ex3*^ adult viability were quantified by calculating observed vs. expected (o/e) Mendelian ratios. A chi-square test was used to analyze significance between samples. Sample sizes (n) and significance *p*-values (p; denoted by asterisks *) are indicated in the text. Graphs and statistical tests were generated using Prism™ (GraphPad Software), with significance level set to p<0.05.

### DATA AVAILABILITY

All *Drosophila* transgenic and mutant lines used in this study are freely available upon request.

## RESULTS

### *Wg/PCP* ALLELES DOMINANTLY MODIFY THE *GMR-Nab2* EYE PHENOTYPE

As described in prior work (Pak *et al*. 2011), overexpression of *Nab2* posterior to the morphogenetic furrow of 3^rd^ instar (L3) eye imaginal discs (*GMR-Nab2*) leads to adult eyes that are rough, reduced in size, and lack red pigmentation in posterior domains (compare **Fig. 1A** to **1B**). Genetic modification of this easily scored phenotype has proven effective in identifying factors that interact functionally with Nab2, including the nuclear poly(A) binding protein Pabp2 (**Fig. 1C**) and the poly(A) polymerase Hiiragi (*hrg*; **Fig. 1D)** (Pak *et al*. 2011) as well as the disease-associated RNA-binding protein Fmr1 (Bienkowski *et al*. 2017). To apply this approach to PCP-Nab2 interactions in a genetic screen, a group of 13 alleles corresponding to core and accessory PCP factors (**Table 1**) were crossed into the *GMR-Nab2* background and scored for modification of eye-roughening and pigment loss in the progeny; only females were scored to avoid hemizygous effects of X-linked genes. As shown in **Table 1**, the 13 alleles can be grouped into four classes based on their phenotypic effects on *GMR-Nab2*: *dsh*^*A3*^, *dsh*^*1*^, *Appl*^*d*^, *puc*^*MI11060*^, *pk*^*pk-sple-14*^, *pk*^*30*^, *tap*^*MI10541*^, *fz*^*JB*^, and *stan*^*frz3*^ alleles act as dominant suppressors of pigment loss and eye roughening (**Fig. 1E-M**); *Wnt4*^*C1*^ acts as a dominant enhancer of pigment loss and eye roughening (**Fig. 1N**); *pygo*^*s123*^ and *vang*^*6*^ have no effect on eye roughening but slightly modify pigment loss; *dsh*^*6*^ and *pk*^*pk-sple-13*^ have no effect on either phenotype (**Fig. 1O** and **Table 1**).

A review of these modifying effects is consistent with a link between Nab2 and the PCP arm of the Wg/PCP pathway. The two *dsh* suppressor alleles *dsh*^*1*^ (K417M) and *dsh*^*A3*^ (R413H) are semi-viable alleles that selectively perturb PCP due to amino acid substitutions in the DEP domain (<u>D</u>ishevelled, <u>E</u>gl-10, <u>P</u>leckstrin) that block Fz-induced translocation of Dsh to the membrane (Axelrod *et al*. 1998; Boutros *et al*. 1998; Penton *et al*. 2002). In contrast, the non-modifying *dsh*^*6*^ allele (also *dsh*^*M20*^) is a lethal amorph (null) that inactivates both classic Wg signaling and the PCP pathway (Perrimon and Mahowald 1987). This pattern of *dsh*^*1*^, *dsh*^*A3*^ vs. *dsh*^*6*^ modification is thus consistent with selective reduction of PCP activity resulting in suppression of *Nab2* overexpression. The suppressor allele *Appl*^*d*^ (*ß-*<u>*A*</u>*myloid* <u>*p*</u>*rotein* <u>*p*</u>*recursor-like*) is an amorph that deletes the central coding region of a transmembrane protein that acts as a PCP-accessory factor in neurons and has established roles in retinal axon pathfinding and synapse formation (Luo *et al*. 1992; Ashley *et al*. 2005; Mora *et al*. 2013; Soldano *et al*. 2013). Two of three alleles of the PCP component *prickle, pk*^*pk-sple14*^ and *pk*^*30*^ (also *Df(2R)pk*) (Gubb *et al*. 1999), also suppress *GMR-Nab2*. One loss-of-function allele each of the Wg/Wnt receptor *frizzled* (*fz*^*JB*^) and the transmembrane PCP component *starry night* (*stan*^*frz3*^; *stan* is also referred to as *flamingo*) also suppresses, while an allele of the Wg/PCP factor *Van Gogh* (*vang*^*6*^) does not. The hypomorphic allele *Wnt4*^*C1*^, which encodes a Fz ligand with roles in canonical and non-canonical Wg/Wnt signaling (Cohen *et al*. 2002), scores as the lone *GMR-Nab2* enhancer. This effect of *Wnt4*^*C1*^ is the opposite of effect of *fz*^*JB*^, which may reflect a proposed antagonistic relationship between these factors in the *Drosophila* eye (Lim *et al*. 2005). Viable P-element insertions into *puckered* (*puc*^*MI11060*^) and *target of Poxn* (*tap*^*MI10541*^) that respectively encode a component of the PCP-regulated JNK pathway (Boutros *et al*. 1998; Martin-Blanco *et al*. 1998) and a regulator of Dsh levels in brain mushroom body (MB) neurons (Yuan *et al*. 2016), also act as *GMR-Nab2* suppressors. Notably, an amorphic allele of *pygopus* (*pygo*^*s123*^), which encodes a key nuclear element of the canonical Wg pathway (Belenkaya *et al*. 2002; Kramps *et al*. 2002; Parker *et al*. 2002), has little effect on the *GMR-Nab2* phenotype. In sum, these data reveal a pattern of genetic interaction between *Nab2* and Wg/PCP alleles in which reduced expression of proteins that act within the PCP-specific arm of the Wg/PCP pathway components mitigate the effect of Nab2 overexpression in the developing eye.

### PCP ALLELES INTERACT WITH ENDOGENOUS *Nab2*

Genetic interactions between multiple PCP alleles and the *GMR-Nab2* overexpression transgene prompted analysis of genetic links between PCP alleles and the genomic allele *Nab2*^*ex3*^, which is a recessive amorph that causes defects in survival, lifespan, locomotion, thoracic bristle morphology and neurodevelopment (Pak *et al*. 2011; Kelly *et al*. 2016). Four different modifiers from the *GMR-Nab2* eye screen – three suppressors, *dsh*^*1*^, *Appl*^*d*^, *pk*^*pk-sple-14*^, and one enhancer, *Wnt4*^*C1*^ - were tested for dominant effects on two easily screened *Nab2*^*ex3*^ null phenotypes: reduced adult survival to eclosion (<5% in past studies e.g. Pak *et al*. 2011), and thoracic bristle kinking. The former *Nab2*^*ex3*^ survival defect was chosen because it can be rescued by neuron-specific expression of *UAS-Nab2* or *UAS-ZC3H14* transgenes (Kelly *et al*. 2016), suggesting that it is an indicator of underlying neuronal function. The latter *Nab2*^*ex3*^ thoracic bristle defect was chosen because it suggests a link between Nab2 and the PCP-regulated process of F-actin assembly (reviewed Yang and Mlodzik 2015).

To analyze adult viability, the percent of pupae hatching into viable adults was calculated for *Nab2*^*pex41*^ (i.e. *wildtype*) control animals and for non-Tubby pupae collected from a *Nab2*^*ex3*^*/TM6B*^*Tb,Hu*^ stock. Consistent with prior work, approximately 3% of *Nab2*^*ex3*^ zygotic nulls hatch as viable adults, while the *Nab2*^*pex41*^ (i.e. *Nab2*^*wt*^) control chromosome confers essentially full viability (97% hatching) (**Fig. 2**). The *dsh*^*1*^ and *Appl*^*d*^ alleles respectively rescue survival to ∼30% and ∼25% among *Nab2*^*ex3*^ females that inherit the *dsh*^*1*^ or *Appl*^*d*^ alleles from hemizygous fathers. The *pk*^*pk-sple14*^ and *Wnt4*^*c1*^ alleles have no significant effect on *Nab2*^*ex3*^ survival. These genetic data show that a subgroup of PCP alleles that modify *GMR-Nab2* also dominantly rescue low viability of *Nab2*^*ex3*^ animals. In *dsh*^*1/+*^;;*Nab2*^*ex3/ex3*^ surviving adult females, this rescue is also associated with reduced thoracic bristle kinking compared to *Nab2*^*ex3*^ null females (∼45% vs. ∼90%; calculated as fraction of individuals showing at least one kinked humeral or scutellar bristle) (**Fig. 3A-B**). These data led us to examine whether loss of *Nab2* is sufficient to produce PCP-like defects in sensitive tissues e.g. altering proximal-distal wing hair orientation, which is a phenotype associated with mutations in many *Drosophila* PCP components (reviewed in Yang and Mlodzik 2015). For this analysis, hairs in a fixed region between L3-L4 veins and distal to the posterior cross vein (PCV) were imaged in control (*Nab2*^*pex41*^; **Fig. 3C**) and *Nab2*^*ex3*^ adult females (**Fig. 3D**). Three examples of adult wing hair morphology in the L3-L4 region are provided for each genotype, together with corresponding magnified insets. Wing hairs in control wings are arrayed in a uniform right-to-left orientation that matches the proximal-to-distal axis of the wing (**Fig. 3C**); by contrast, hairs in *Nab2*^*ex3*^ wings show consistent orientation defects (**Fig. 3D**), including misrotation of individual hairs and occasional evidence of coordinated misrotation among adjacent hair cells (e.g. **Fig. 3D**, inset in top panel). These PCP-like hair defects in *Nab2* null females are consistent with a role for Nab2 in mechanisms that guide planar polarity of wing cells.

**Figure 3.**
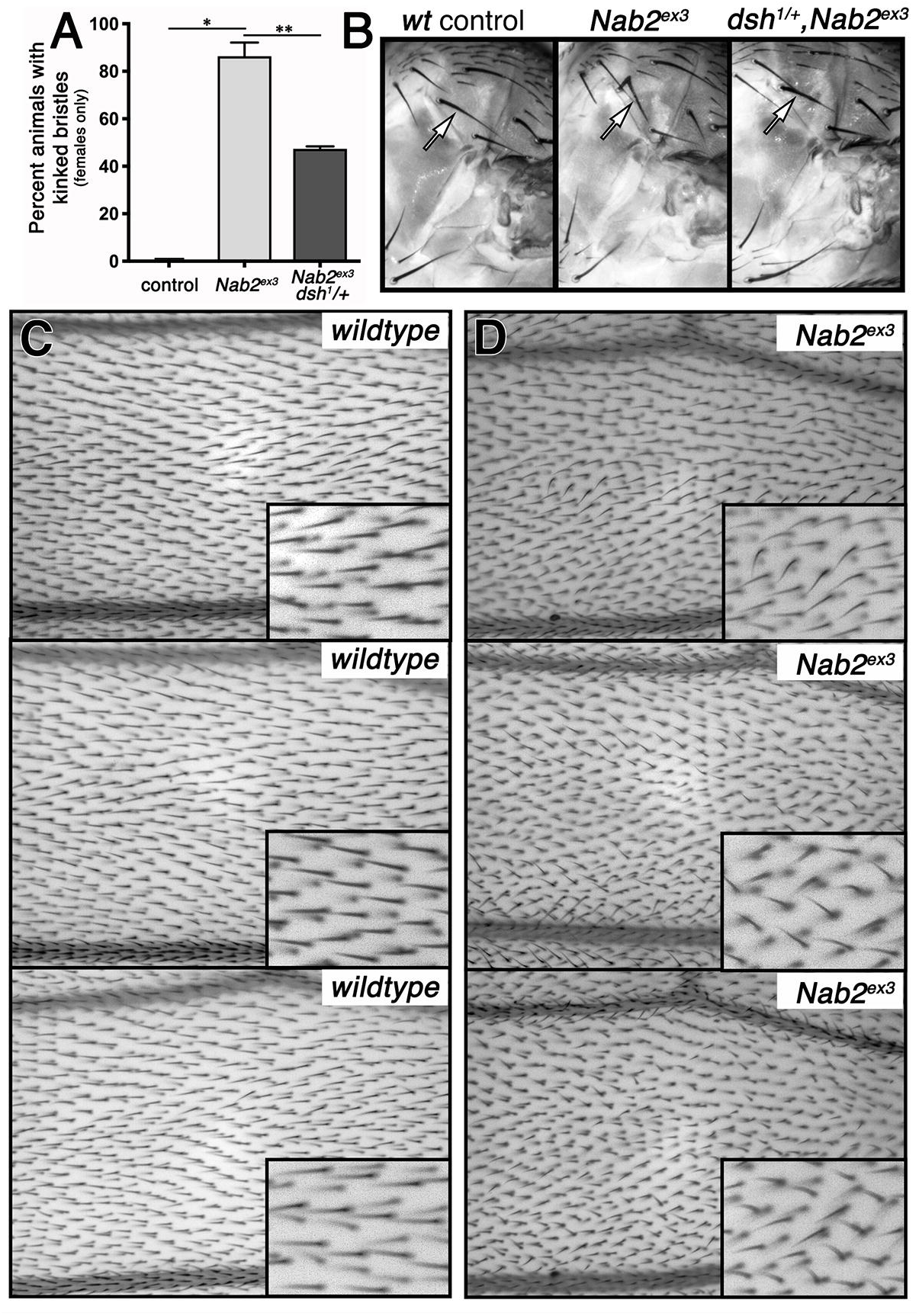
*Nab2*^*ex3*^ effects on bristle morphology and wing hair orientation. (**A**) Frequency of bristle kinking in *Nab2*^*pex41*^ (control), *Nab2*^*ex3*^, and *dsh*^*1*^*/+;;Nab2*^*ex3*^ adult females (n=3 separate crosses). Note suppression by *dsh*^*1*^ heterozygosity. Statistical significance is indicated (* p=0.0003; **p=0.02). (**B**) Examples of humeral bristle morphology (arrows) in the same genotypes as in **A**. Images of wing hairs in the L3-L4 region from three representative examples of (**C**) control (*Nab2*^*pex41*^) or (**D**) *Nab2*^*ex3*^ adult female wings orientated proximal to distal (right to left). Insets (lower right) show magnified views from each panel.

### PCP AND *Nab2* GENETIC INTERACTIONS THROUGH THE PATERNAL GERMLINE

In the course of analyzing modified *GMR-Nab2* progeny descended from *dsh*^*1*^ and *Appl*^*d*^ hemizygous fathers (i.e. *dsh*^*1*^*/Y* and *Appl*^*d*^*/Y*), we noted that male offspring carrying the *GMR-Nab2* transgene exhibit suppression of eye roughness that approaches that seen in their *dsh*^*1*^*/+* and *Appl*^*d*^*/+* heterozygous sisters (compare **Fig. 4C,D** vs. **Fig. 1F,G**). This result is unexpected because these males inherit a paternal *Y* chromosome, rather than *dsh*^*1*^ and *Appl*^*d*^ alleles, and because the unmodified *GMR-Nab2* phenotype is more severe in males than females (see Fig. 1B vs. 4B and Pak *et al*. 2011). This observed suppression through the male germline was not due inheritance of an X-chromosome balancer and could not be replicated by direct RNAi depletion of *dsh* in the germline (data not shown; *nanos-Gal4,UAS-dsh-RNAi*, BDSC lines #31306 and #31307). The indirect effect of *dsh*^*1*^ and *Appl*^*d*^ alleles on *GMR-Nab2* might be explained by heritable changes in the *dsh*^*1*^ and *Appl*^*d*^ hemizygous male germline that are passed to progeny. Germline roles for *Drosophila* PCP components have not been reported; however, the *C. elegans vang* homolog VANG-1 acts upstream of the insulin/IGF-1 responsive DAF-16/FoxO transcription factor in the germline to control lifespan (Honnen *et al*. 2012). This PCP-to-nucleus link has not been explored in the *Drosophila* germline, but it could provide a mechanism through which PCP alleles generate epigenetic changes in the male germline that modify - *GMR-Nab2* phenotypes in progeny. Importantly, we also observed evidence of this type of link between endogenous *Nab2* and *dsh*. A paternal copy of *dsh*^*1*^ indirectly rescues survival among *+/Y;;Nab2*^*ex3/ex3*^ male offspring to ∼23% (**Fig. 1E**), which mirrors the indirect *dsh*^*1*^ rescue of *GMR-Nab2* eye roughening. By contrast, the *Appl*^*d*^ allele has no indirect effect on the survival of *+/Y;;Nab2*^*ex3/ex3*^ male progeny. This difference between the paternal effects of *dsh*^*1*^ and *Appl*^*d*^ on *+/Y;;Nab2*^*ex3/ex3*^ offspring is consistent with a strong germline link between *Nab2* and *dsh*. In sum, these rescue data imply two modes of *Nab2* null rescue by PCP loss-of-function alleles: (1) by direct zygotic inheritance of *dsh*^*1*^ or *Appl*^*d*^, which could suggest that Nab2 normally represses PCP activity; (2) and by indirect exposure to *dsh*^*1*^, but not *Appl*^*d*^, in the male germline.

**Figure 4.**
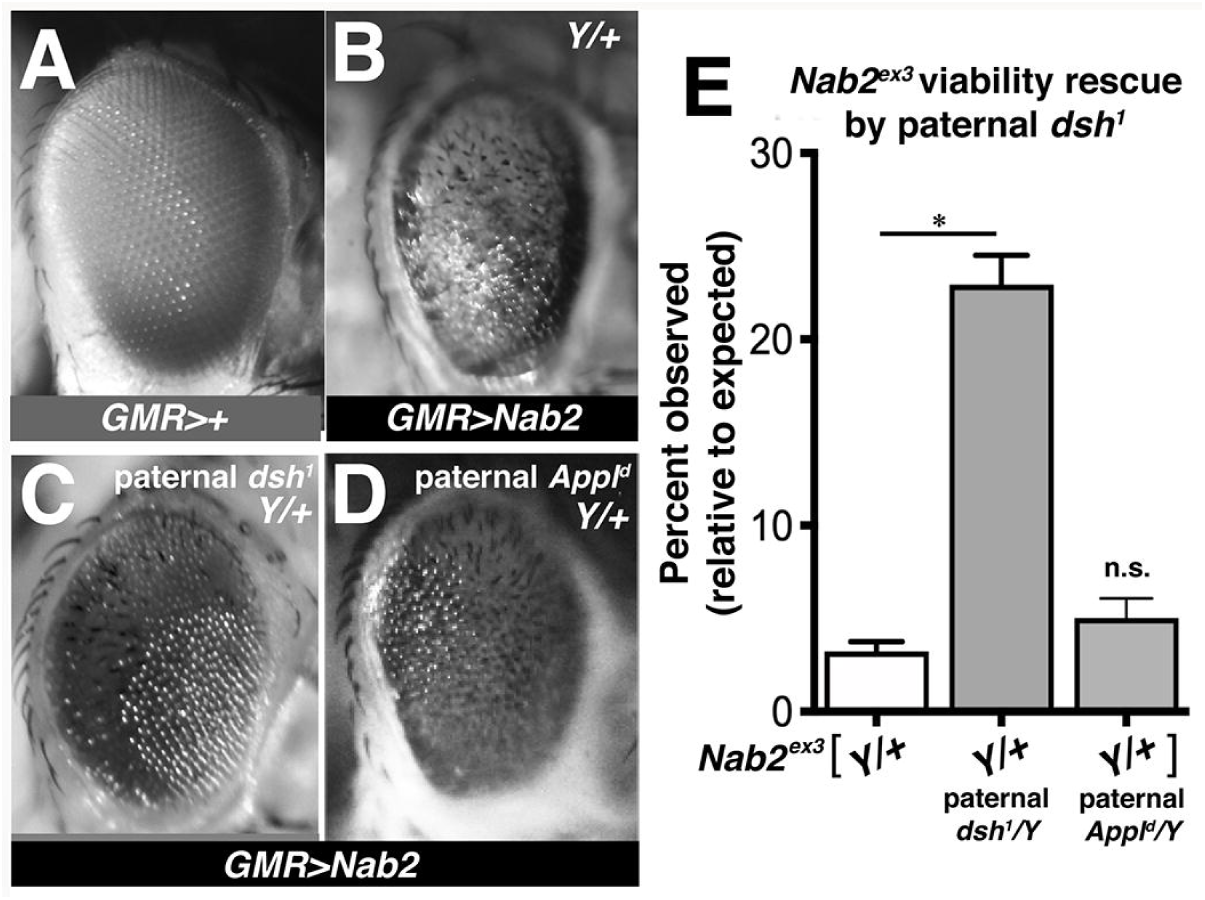
A *Nab2-dsh* genetic interaction through the male germline. Images of adult male eyes from (**A**) control, (**B**) *GMR-Nab2*, or *Y/+,GMR-Nab2* progeny of (**C**) *dsh*^*1*^*/Y* or (**D**) *Appl*^*d*^*/Y* fathers. (**E**) Histogram of eclosion frequencies of unmodified male *Nab2*^*ex3*^ flies, or the *Y/+,Nab2*^*ex3/ex3*^ progeny of *dsh*^*1*^*/Y* or *Appl*^*d*^*/Y* fathers (respectively labelled “*Y/+*”, “*Y/+ paternal dsh*^*1*^*/Y*”, and “*Y/+ paternal Appl*^*d*^*/Y*”). Statistical significance is indicated (* p=0.0008; n.s.= not significant).

## DISCUSSION

Here we have used three phenotypes caused by altered dosage of the *Drosophila* poly(A) RNA-binding protein Nab2, (1) eye roughness caused by overexpression of *Nab2* (*GMR-Nab2*), (2) neuronal requirement for *Nab2* in adult survival (Kelly *et al*. 2014) and (3) thoracic bristle kinking in *Nab2* null animals, to screen for genetic interactions between *Nab2* and genes encoding components of the Wg/planar cell polarity (PCP) pathway. The *Nab2*-PCP interactions detected by this approach are consistent with a role for Nab2 in restraining PCP signaling in certain <u>*Drosophila* tissues</u>, perhaps by inhibiting expression of a PCP component *in vivo*. As Nab2 and its human ortholog ZC3H14 play conserved roles in neurodevelopment and neural function (Pak *et al*. 2011; Kelly *et al*. 2014; Kelly *et al*. 2016; Bienkowski *et al*. 2017; Rha *et al*. 2017; Collins *et al*. 2019), these data provide insight into pathways that may be disrupted in ZC3H14-associated intellectual disability.

The suppressive effects of alleles that impair Wg/PCP signaling, and in some cases specifically perturb the PCP arm of the Wg pathway (e.g. *dsh*^*1*^, *Appl*^*d*^), are observed with phenotypes caused by both *Nab2* overexpression (*GMR-Nab2*) and loss (*Nab2*^*ex3/ex3*^). This pattern differs from other *Nab2* modifier alleles that have inverse effects in these two *Nab2* backgrounds (for example *dfmr1*^Δ*50*^ as in Bienkowski *et al*. 2017). The common suppressive effect of *dsh*^*1*^ and *Appl*^*d*^ on *Nab2* gain/loss phenotypes could be explained if a normal dose of Nab2 is required to maintain a balance of PCP signaling between cells, and that imbalanced PCP signaling resulting from Nab2 overexpression or loss is then restored by reducing the genetic dose of either *dsh, Appl, puc, pk, tap, fz*, or *stan*.

Among Wg/PCP alleles tested in this study, the *dsh*^*1*^ allele displayed the most consistent suppressive effects on *Nab2* null phenotypes. Because *dsh*^*1*^ specifically impairs PCP signaling (Axelrod *et al*. 1998), *Nab2* null flies might thus be expected to display defects in hallmark PCP-regulated processes, e.g. wing hair polarization and ommatidial rotation. Consistent with this hypothesis, we demonstrate here that *Nab2* loss causes hair misorientation in adult wings, suggesting that Nab2 regulates PCP within wing cells. As Nab2 is expressed ubiquitously but required in neurons for viability (Kelly *et al*. 2014), the rescue of *Nab2* null viability by *dsh*^*1*^ implies that the Nab2-PCP link may also occur in neurons. This idea is further supported by the genetic interaction between *Nab2*^*ex3*^ and the *Appl*^*d*^ allele, which inactivates a neuron-specific PCP component (Soldano *et al*. 2013), and by *GMR-Nab2* modification by an allele of *tap*, a regulator of *dsh* expression in neurons (Yuan *et al*. 2016). Interestingly, both *Nab2* and PCP alleles individually alter the trajectories of axons that project from a group of brain neurons termed Kenyon cells (Ng 2012; Kelly *et al*. 2016), which provides a cellular context for future study of the Nab2-PCP interaction.

Nab2 may modify PCP-regulated developmental processes indirectly through post-transcriptional effects that regulate expression of a protein(s) that operates in parallel to PCP (e.g. via F-actin bundling). Alternatively, Nab2 may directly regulate post-transcriptional expression of a PCP component *in vivo*. Work on the Nab2 mammalian homolog ZC3H14 homolog has identified PCP factors whose mRNAs are candidate Nab2/ZC3H14 targets. The hippocampal proteome of *Zc3h14* knockout mice contains elevated levels of the Vang family member Vangl2 (Rha *et al*. 2017), which is a component of the vertebrate PCP pathway (reviewed in Bailly *et al*. 2018). A separate study found that ZC3H14 depletion leads to intron retention in the *PSD95* mRNA (Morris and Corbett 2018), which encodes a postsynaptic guanylate kinase required for activity-dependent synaptic plasticity (reviewed in Xu 2011). Intriguingly, the fly PSD95 homolog Discs Large-1 (Dlg) controls canonical Wg signaling in wing tissue by stabilizing Dsh protein (Liu *et al*. 2016). In light of these observations, the corresponding fly *vang* and *dlg1* mRNAs are candidate Nab2 targets that may contribute to Nab2-PCP genetic interactions documented in this study.

In the course of these studies, we found evidence of an unexpected link between PCP and *Nab2* in the male germline. Hemizygosity for *dsh*^*1*^ in the paternal germline can rescue eye roughening in *+/Y;GMR-Nab2* males and survival of *+/Y;;Nab2*^*ex3/ex3*^ males, while *Appl*^*d*^ hemizygosity in the paternal germline rescues eye roughening in *+/Y;GMR-Nab2* males but not *+/Y;;Nab2*^*ex3/ex3*^ survival. Maternal effect mutants are quite common and reflect the important role of maternally provisioned mRNAs and proteins in early development (Schupbach and Wieschaus 1986). Paternal effects are by comparison much less common and have not been reported previously for *dsh* or *Appl* alleles. One potential explanation for the *dsh*^*1*^/*Appl*^*d*^ hemizygous effects could be that these alleles produce heritable epigenetic changes in the paternal genome that affect expression of factors involved in *GMR-Nab2* and/or *Nab2*^*ex3*^ phenotypes in the subsequent generation. However, the PCP pathway is generally regarded as a cytoplasmic circuit that locally remodels the cytoskeleton but lacks nuclear output (e.g. as in the *Drosophila* wing). Notably, one study suggests that signals in the *C. elegans* germline from the Vang homolog VANG1 may be transmitted into the nucleus by the DAF-16/FoxO transcription factor (Honnen *et al*. 2012). This finding links a core PCP component, VANG-1, to the activity of a transcription factor that could alter heritable chromatin states. The analogous Vang-FoxO link has not been studied in the fly germline but it is a candidate to mediate the paternal effect of *dsh*^*1*^ and *Appl*^*d*^ alleles on *Nab2* alleles.

In sum, here we present the results of a candidate-based genetic screen that identifies a series of dominant genetic interactions between Wg/PCP alleles and both a *Nab2* transgene and null allele. Collectively these interactions provide evidence that the Nab2 RNA binding protein may regulate expression of PCP component(s) in different cell types, including neurons and/or wing hair cells. This latter conclusion is significant given that very few Nab2-target mRNAs are known and that the Nab2 human ortholog ZC3H14 is lost in an inherited form of recessive intellectual disability (Pak *et al*. 2011). Identifying a conserved link between Nab2/ZC3H14 and PCP activity would be a significant step toward better understanding the conserved role of these proteins in development and disease. Based on the strength of the interactions between *Nab2* and PCP alleles, our data can now be used to generate and test hypotheses of how the Nab2 RNA binding protein is physically linked to PCP components *in vivo*.

## ACKNOWLEDGMENTS

Stocks obtained from the Bloomington Drosophila Stock Center (NIH P40OD018537) were used in this study. We thank members of the Moberg, Corbett, Chen and Caspary labs for helpful discussion, and G. Pierre-Louis for sharing stocks collected from the Axelrod Lab (Stanford). This work was funded by the National Institute of Health MH10730501 to K.H.M. and A.H.C.

## LEGENDS

**Table 1. Summary of tested alleles and their effect of *GMR-Nab2* morphology**. Modification of pigment loss and overall eye structure in *GMR-Nab2* adult females (*GMR-Gal4/+;Nab2*^*EP3716*^*/+*) was scored separately for each of the thirteen PCP alleles tested indicated (“Sup” = suppressed, “Enh” = enhanced).

## Bibliography

Ackley, B. D., 2014 Wnt-signaling and planar cell polarity genes regulate axon guidance along the anteroposterior axis in C. elegans. Dev Neurobiol 74: 781–796.

Ashley, J., M. Packard, B. Ataman and V. Budnik, 2005 Fasciclin II signals new synapse formation through amyloid precursor protein and the scaffolding protein dX11/Mint. J Neurosci 25: 5943–5955.

Aviles, E. C., and E. T. Stoeckli, 2016 Canonical wnt signaling is required for commissural axon guidance. Dev Neurobiol 76: 190–208.

Axelrod, J. D., J. R. Miller, J. M. Shulman, R. T. Moon and N. Perrimon, 1998 Differential recruitment of Dishevelled provides signaling specificity in the planar cell polarity and Wingless signaling pathways. Genes Dev 12: 2610–2622.

Bailly, E., A. Walton and J. P. Borg, 2018 The planar cell polarity Vangl2 protein: From genetics to cellular and molecular functions. Semin Cell Dev Biol 81: 62–70.

Belenkaya, T. Y., C. Han, H. J. Standley, X. Lin, D. W. Houston et al., 2002 pygopus Encodes a nuclear protein essential for wingless/Wnt signaling. Development 129: 4089–4101.

Bienkowski, R. S., A. Banerjee, J. C. Rounds, J. Rha, O. F. Omotade et al., 2017 The Conserved, Disease-Associated RNA Binding Protein dNab2 Interacts with the Fragile X Protein Ortholog in Drosophila Neurons. Cell Rep 20: 1372–1384.

Boutros, M., N. Paricio, D. I. Strutt and M. Mlodzik, 1998 Dishevelled activates JNK and discriminates between JNK pathways in planar polarity and wingless signaling. Cell 94: 109–118.

Castello, A., B. Fischer, M. W. Hentze and T. Preiss, 2013 RNA-binding proteins in Mendelian disease. Trends Genet 29: 318–327.

Cohen, E. D., M. C. Mariol, R. M. Wallace, J. Weyers, Y. G. Kamberov et al., 2002 DWnt4 regulates cell movement and focal adhesion kinase during Drosophila ovarian morphogenesis. Dev Cell 2: 437–448.

Collins, S. C., A. Mikhaleva, K. Vrcelj, V. E. Vancollie, C. Wagner et al., 2019 Large-scale neuroanatomical study uncovers 198 gene associations in mouse brain morphogenesis. Nat Commun 10: 3465.

Fasken, M. B., A. H. Corbett and M. Stewart, 2019 Structure-function relationships in the Nab2 polyadenosine-RNA binding Zn finger protein family. Protein Sci 28: 513–523.

Gombos, R., E. Migh, O. Antal, A. Mukherjee, A. Jenny et al., 2015 The Formin DAAM Functions as Molecular Effector of the Planar Cell Polarity Pathway during Axonal Development in Drosophila. J Neurosci 35: 10154–10167.

Gubb, D., C. Green, D. Huen, D. Coulson, G. Johnson et al., 1999 The balance between isoforms of the prickle LIM domain protein is critical for planar polarity in Drosophila imaginal discs. Genes Dev 13: 2315–2327.

Hollis, E. R., 2nd, and Y. Zou, 2012 Expression of the Wnt signaling system in central nervous system axon guidance and regeneration. Front Mol Neurosci 5: 5.

Honnen, S. J., C. Büchter, V. Schröder, M. Hoffmann, Y. Kohara et al., 2012 C. elegans VANG-1 Modulates Life Span via Insulin/IGF-1-Like Signaling. PLoS ONE 7.

Inaki, M., S. Yoshikawa, J. B. Thomas, H. Aburatani and A. Nose, 2007 Wnt4 is a local repulsive cue that determines synaptic target specificity. Curr Biol 17: 1574–1579.

Kelly, S. M., R. Bienkowski, A. Banerjee, D. J. Melicharek, Z. A. Brewer et al., 2016 The Drosophila ortholog of the Zc3h14 RNA binding protein acts within neurons to pattern axon projection in the developing brain. Dev Neurobiol 76: 93–106.

Kelly, S. M., S. W. Leung, C. Pak, A. Banerjee, K. H. Moberg et al., 2014 A conserved role for the zinc finger polyadenosine RNA binding protein, ZC3H14, in control of poly(A) tail length. RNA 20: 681–688.

Kramps, T., O. Peter, E. Brunner, D. Nellen, B. Froesch et al., 2002 Wnt/wingless signaling requires BCL9/legless-mediated recruitment of pygopus to the nuclear beta-catenin-TCF complex. Cell 109: 47–60.

Lim, J., K. K. Norga, Z. Chen and K. W. Choi, 2005 Control of planar cell polarity by interaction of DWnt4 and four-jointed. Genesis 42: 150–161.

Liu, M., Y. Li, A. Liu, R. Li, Y. Su et al., 2016 The exon junction complex regulates the splicing of cell polarity gene dlg1 to control Wingless signaling in development. Elife 5.

Luo, L., T. Tully and K. White, 1992 Human amyloid precursor protein ameliorates behavioral deficit of flies deleted for Appl gene. Neuron 9: 595–605.

Martin-Blanco, E., A. Gampel, J. Ring, K. Virdee, N. Kirov et al., 1998 puckered encodes a phosphatase that mediates a feedback loop regulating JNK activity during dorsal closure in Drosophila. Genes Dev 12: 557–570.

Mora, N., I. Almudi, B. Alsina, M. Corominas and F. Serras, 2013 beta amyloid protein precursor-like (Appl) is a Ras1/MAPK-regulated gene required for axonal targeting in Drosophila photoreceptor neurons. J Cell Sci 126: 53–59.

Morris, K. J., and A. H. Corbett, 2018 The polyadenosine RNA-binding protein ZC3H14 interacts with the THO complex and coordinately regulates the processing of neuronal transcripts. Nucleic Acids Res 46: 6561–6575.

Ng, J., 2012 Wnt/PCP proteins regulate stereotyped axon branch extension in Drosophila. Development 139: 165–177.

Pak, C., M. Garshasbi, K. Kahrizi, C. Gross, L. H. Apponi et al., 2011 Mutation of the conserved polyadenosine RNA binding protein, ZC3H14/dNab2, impairs neural function in Drosophila and humans. Proc Natl Acad Sci U S A 108: 12390–12395.

Parker, D. S., J. Jemison and K. M. Cadigan, 2002 Pygopus, a nuclear PHD-finger protein required for Wingless signaling in Drosophila. Development 129: 2565–2576.

Penton, A., A. Wodarz and R. Nusse, 2002 A mutational analysis of dishevelled in Drosophila defines novel domains in the dishevelled protein as well as novel suppressing alleles of axin. Genetics 161: 747–762.

Perrimon, N., and A. P. Mahowald, 1987 Multiple functions of segment polarity genes in Drosophila. Dev Biol 119: 587–600.

Rha, J., S. K. Jones, J. Fidler, A. Banerjee, S. W. Leung et al., 2017 The RNA-binding Protein, ZC3H14, is Required for Proper Poly(A) Tail Length Control, Expression of Synaptic Proteins, and Brain Function in Mice. Hum Mol Genet.

Sato, M., D. Umetsu, S. Murakami, T. Yasugi and T. Tabata, 2006 DWnt4 regulates the dorsoventral specificity of retinal projections in the Drosophila melanogaster visual system. Nat Neurosci 9: 67–75.

Schupbach, T., and E. Wieschaus, 1986 Germline autonomy of maternal-effect mutations altering the embryonic body pattern of Drosophila. Dev Biol 113: 443–448.

Soldano, A., Z. Okray, P. Janovska, K. Tmejova, E. Reynaud et al., 2013 The Drosophila homologue of the amyloid precursor protein is a conserved modulator of Wnt PCP signaling. PLoS Biol 11: e1001562.

Srahna, M., M. Leyssen, C. M. Choi, L. G. Fradkin, J. N. Noordermeer et al., 2006 A signaling network for patterning of neuronal connectivity in the Drosophila brain. PLoS Biol 4: e348.

Vandewalle, J., M. Langen, M. Zschatzsch, B. Nijhof, J. M. Kramer et al., 2013 Ubiquitin ligase HUWE1 regulates axon branching through the Wnt/beta-catenin pathway in a Drosophila model for intellectual disability. PLoS One 8: e81791.

Xu, W., 2011 PSD-95-like membrane associated guanylate kinases (PSD-MAGUKs) and synaptic plasticity. Curr Opin Neurobiol 21: 306–312.

Yang, Y., and M. Mlodzik, 2015 Wnt-Frizzled/planar cell polarity signaling: cellular orientation by facing the wind (Wnt). Annu Rev Cell Dev Biol 31: 623–646.

Yuan, L., S. Hu, Z. Okray, X. Ren, N. De Geest et al., 2016 The Drosophila neurogenin Tap functionally interacts with the Wnt-PCP pathway to regulate neuronal extension and guidance. Development 143: 2760–2766.

